# Mechanism study on lactic acid promoting intervertebral disc degeneration by regulating senescence and DNA damage of cartilage endplate stem cells

**DOI:** 10.64898/2026.01.23.701236

**Authors:** Qiao Lv, Tianling Wang, Lu Jiang, Qian Chen, Jin Peng, Ju Zhou, Qian Min, Yongqi Pu, Jianyun Zhou, Qing Huang

**Author notes:** Clinical Research Center, Xinqiao Hospital, Army Medical University, Chongqing 400037, China. These authors contributed equally to this work (Qiao Lv, Tianling Wang).

## Abstract

Intervertebral disc degeneration, a leading cause of low back pain with incompletely elucidated molecular mechanisms, was studied via integrated in vivo/vitro approaches. This study first reveals that lactic acid accelerates intervertebral disc degeneration by inducing cartilage endplate stem cells senescence and DNA damage, thereby activating the P16/P21/P53-centered senescence pathway. In a rat tail vertebra puncture-induced intervertebral disc degeneration model, degenerated discs exhibited increased lactic acid levels, narrowed intervertebral spaces, and disrupted nucleus pulposus structure (*P*<0.05). In vitro, 0/2/6/10 mM lactic acid dose-dependently suppressed cartilage endplate stem cells viability (10 mM group: 15.7% of the control), elevated intracellular reactive oxygen species (ROS, 2.8-fold relative to the control), induced G0 cell cycle arrest (10 mM group: 85.63%), reduced EdU-positive cells (8.62%), and increased β-galactosidase-positive cells (10 mM group: 33.06%) and γ-H2AX foci (all *P*<0.01).Molecularly, lactic acid significantly upregulated P16 (2.1-fold), P21 (3.1-fold), P53 (2.4-fold), and γ-H2AX (1.8-fold). In vivo intervertebral disc injection confirmed a positive correlation between lactic acid concentration and intervertebral disc degeneration severity. This study clarifies lactic acid’s role in intervertebral disc degeneration via the “oxidative stress–cell cycle arrest–cellular senescence” axis, advancing understanding of intervertebral disc degeneration pathogenesis and providing a basis for targeted therapies against lactic acid metabolism.

## Introduction

Intervertebral disc degeneration (IDD), a key pathological driver of low back pain and a range of spine-related diseases, not only severely compromises patients’ physical health but also substantially impacts their daily activity capacity, mental state, and overall quality of life. It has thus emerged as one of the critical pathological foundations of spinal diseases that demand prioritized attention in clinical practice^1–5^. With the continuous acceleration of population aging globally, particularly in China, the incidence rate of IDD has exhibited a steady annual upward trend. This epidemiological trend not only exposes more people to the distress of symptoms like low back pain but also imposes mounting challenges on the clinical prevention and treatment of spinal diseases ^6,7^. The typical pathological features of this disease are characterized by nucleus pulposus dehydration, annulus fibrosus rupture, and cartilage endplate calcification^8,9^, these structural changes gradually undermine the biomechanical stability of the intervertebral disc, thereby inducing relevant clinical symptoms. While existing studies have confirmed that multiple factors—including hypoxic microenvironment, chronic inflammatory response, and abnormal mechanical stress—are involved in the pathological progression of IDD ^10–12^, the core driving mechanisms regulating the disease’s development remain incompletely elucidated. Thus, there is an urgent need for further in-depth research to enhance the understanding of its pathological underpinnings.

In recent years, the abnormal metabolic microenvironment has been garnering growing attention within the academic community for its regulatory role in the pathological progression of IDD. Analysis of intervertebral disc samples from clinical IDD patients reveals that lactic acid concentration in degenerated disc tissue is significantly elevated, with levels reaching up to 3–5 times those of normal discs; this clinical observation indicates that abnormal accumulation of lactic acid in degenerated discs may be one of the key pathological features in the development of IDD^13–15^. Abnormal lactic acid accumulation is currently thought to be potentially associated with metabolic reprogramming in intervertebral disc cells^16–18^. However, the specific regulatory role lactic acid exerts in IDD development, along with the underlying molecular mechanisms, have yet to be systematically elucidated. This has also emerged as one of the key issues in urgent need of breakthroughs in current research into IDD’s pathological mechanisms.

Cartilage endplate stem cells (CESCs) reside at the core interface for nutrient exchange in the intervertebral disc, a location enabling them to directly contribute to regulating the disc microenvironment. As an important cell population with repair potential in the intervertebral disc, CESCs not only serve the core function of maintaining disc matrix homeostasis but also exhibit the key potential to mediate repair after disc injury—making them crucial for the structural integrity and functional maintenance of the disc^19^. If CESCs exhibit dysfunction, they can directly trigger an imbalance between the synthesis and degradation processes of the extracellular matrix (ECM)^19,20^; This imbalance further disrupts the dynamic stability of the disc matrix, leading to sustained damage to the disc’s structure and function and ultimately significantly accelerating the pathological progression of IDD. Notably, degenerated intervertebral disc tissue exhibits a characteristic acidic microenvironment (with pH values ranging from 6.5 to 6.8), and lactic acid is the key contributing factor to this acidic microenvironment^21^. This specific acidic microenvironment is highly likely to influence the pathological progression of IDD by precisely modulating the core biological behaviors of CESCs, including cell proliferation, directed differentiation, and senescence processes^22,23^.

The present study seeks to systematically elucidate the biological action mechanism of lactic acid in IDD. By establishing a classical needle puncture-induced rat caudal intervertebral disc degeneration model and combining it with *in vitro* and *in vivo* functional experiments of lactic acid intervention, this study focuses on exploring the regulatory effects of lactic acid on cartilage endplate stem cell (CESC) functions—particularly the specific impacts on the senescent phenotype and DNA damage of CESCs. It further reveals the core pathological links through which lactic acid drives intervertebral disc degeneration via regulating CESC functions, ultimately providing a novel theoretical basis and research insights for developing precision-targeted therapeutic strategies for IDD.

## Materials and Methods

### 1. Animal experiments

The animal experiments in this study was approved by the Laboratory Animal Welfare and Ethics Committee of the Army Medical University, PLA(NO AMUWE20255413). Animals were randomly assigned to groups, and all experiments were performed according to standard laboratory procedures of randomization and blinding. Rats(2-month-old male Sprague-Dawley) in the normal group received no treatment, while those in the PIDD group were subjected to procedures based on previous literature^24,25^. Rats were anesthetized with 2% pentobarbital (50 mg/kg) and administered subcutaneous carprofen (5 mg/kg) for analgesia before surgery. After disinfecting the surgical area with iodophor, a 20 g sterile puncture needle was vertically inserted into the Co7/Co8 intervertebral space of the tail vertebrae, with a depth of approximately 5 mm. The needle was rotated 360 degrees, kept in place for 30 seconds, then removed vertically, and the surgical area was disinfected again. Postoperatively, carprofen (5 mg/kg/day) for analgesia and antibiotics for infection prevention were given for 3 consecutive days. Eight weeks after surgery, rats were euthanized after deep anesthesia with sodium pentobarbital. The caudal vertebrae and target intervertebral discs (Co7/Co8) were completely removed for subsequent lactic acid content determination, histological, and radiological analyses.

In in vivo injection experiments, rats were randomly allocated into 4 groups (n = 6–8 per group). 0 mM (control), 2 mM, 6 mM, and 10 mM lactic acid solutions were injected into the Co7/Co8 intervertebral space of the caudal vertebrae at a uniform volume of 5 μL per disc. Four weeks post-injection, intervertebral discs were harvested for subsequent assays.

### 2. Cell isolation, culture, and treatment

According to previous literature^26^, under sterile conditions, endplate cartilage tissue was obtained, minced, and then digested with 0.2% type II collagenase (C2-BIOC, Sigma–Aldrich, USA) at 37°C for approximately 2 hours. The digested product was filtered through a 70 μm cell strainer, centrifuged at 400 × g, and the cell pellet was resuspended in DMEM/F12 medium (BI, Kibbutz Beit-Haemek, Israel) supplemented with 10% fetal bovine serum (FBS; C04001500, VivaCell, Shanghai, China) and 1% penicillin/streptomycin (C0222, Beyotime, Shanghai, China).Cartilage endplate stem cells (CESCs) were cultured in an incubator at 37°C with 5% CO₂. The medium was changed every 3–4 days, and passaging was performed when cell confluency reached 80–90%. The third passage of CESCs was used in all experiments. For lactic acid treatment, CESCs were exposed to lactic acid (L6402, Sigma-Aldrich) at concentrations of 0 mM, 2 mM, 6 mM, and 10 mM for 24 hours. After treatment, the cells were harvested for subsequent assays.

### 3. Analysis of lactic acid content

The lactic acid content in intervertebral disc tissue was measured using a lactic acid assay kit (BC2235, Solarbio, China). Briefly, 50 mg of fresh nucleus pulposus (NP) tissue was added to the extraction solution, followed by thorough homogenization using an ultrasonic homogenizer (M3000, Biologics, USA). The supernatant was then collected. A standard curve was constructed according to the manufacturer’s instructions, after which the reaction mixture was added to standard or sample wells, and the absorbance was measured at 570 nm.The lactic acid concentration of the samples was calculated using the following formula:Lactic acid concentration (nmol/mg) = (LA / SV) × D Where:LA = Lactic acid amount in the well calculated from the standard curve (nmol),SV = Weight of sample added to the well (mg),D = Sample dilution factor.

### 4. Radiological evaluation

Eight weeks after surgery, X-ray imaging of rat tail vertebrae was performed (Bruker SkyScan1176). Referring to the method in the literature^24^, Image J software was used to measure the length of the target intervertebral disc and adjacent upper and lower vertebrae, and the disc height index (DHI) was calculated. The formula for DHI percentage (DHI%) is: DHI% = (DHI at 8 weeks after surgery / DHI before surgery) * 100%.

### 5. Histological analysis

Collected rat tail vertebrae and intervertebral disc samples were fixed in 4% paraformaldehyde solution (Beyotime) for 48 hours, then decalcified with 10% EDTA (pH 7.2) for 14 days. Samples were dehydrated with gradient ethanol, cleared with xylene, and embedded in paraffin. Paraffin blocks were sectioned continuously (3 μm thickness). Sections were stained with hematoxylin-eosin (HE) (G1120, Solarbio) and modified Safranin O-Fast Green (SO & FG) (G1371, Solarbio) respectively. After the stained sections were sealed with neutral gum, they were observed under an optical microscope (BX53, OLYMPUS). The degree of intervertebral disc degeneration was scored according to the published histological grading criteria for intervertebral disc degeneration^27^.

### 6. Cell counting Kit-8 (CCK-8) assay

The effect of lactic acid on the viability of cartilage endplate stem cells (CESCs) was assessed using a Cell Counting Kit-8 (CCK-8; C0038, Beyotime). CESCs were seeded into 96-well plates at a density of 2 × 10³ cells/well and allowed to adhere for 12 hours, followed by treatment with lactic acid at concentrations of 0 mM, 2 mM, 6 mM, and 10 mM, respectively.At 0 h, 12 h, 24 h, 36 h, and 48 h post-treatment, 10 μl of CCK-8 reagent was added to each well. The plates were then incubated at 37°C for 2 hours in the dark, and the absorbance was measured at 450 nm using a microplate reader (M2, Molecular Devices, USA).

### 7. EdU staining assay

The effect of lactic acid on the proliferation of cartilage endplate stem cells (CESCs) was detected using an EdU-555 Cell Proliferation Assay Kit (C0075S, Beyotime). CESCs were seeded into 6-well plates at a density of 2 × 10⁵ cells/well.

Following 24 hours of lactic acid treatment, 2 ml of medium containing 10 μM EdU was added to each well, and incubation was continued for 2 hours. The cells were then fixed with 4% paraformaldehyde for 10 minutes, followed by incubation with immunostaining permeabilization solution (P0097, Beyotime) for 15 minutes. Subsequently, 2 ml of Click reaction solution (prepared according to the kit instructions) was added, and the cells were incubated at room temperature in the dark for 30 minutes. Nuclear staining was performed with Hoechst 33342 for 15 minutes.Finally, the cells were observed and imaged under a fluorescence microscope (IX 73, Olympus, Tokyo, Japan), and the EdU-positive cell rate was calculated.

### 8. Cell cycle assay

PI/RNase staining buffer (550825, BD Biosciences) and FITC-labeled Ki-67 antibody (11882S, CST) were used for cell cycle detection. CESCs were seeded at a density of 4×10⁵ cells/T25 flask. When the cell confluency reached 70%, lactic acid treatment (0-10 mM, 24 h) was performed. Cells were collected, washed with PBS, and fixed with pre-cooled 70% ethanol at 4°C overnight. Fixed cells were washed with PBS and centrifuged at 400×g for 5 min. The supernatant was discarded, and Ki-67 antibody (1:50 diluted in PBS) was added and incubated at room temperature in the dark for 1 hour. After washing with PBS, PI/RNase staining buffer was added and incubated at room temperature in the dark for 15 min. Cell cycle distribution was analyzed using a flow cytometer (Gallios, Beckman Coulter, USA) (Ki-67 vs. PI dual parameters).

### 9. Senescence-associated β-galactosidase (SA-β-gal) staining assay

Senescence-associated β-galactosidase (SA-β-gal) staining kit (C0602, Beyotime) was used to detect lactic acid-induced senescence of CESCs. CESCs were seeded in 6-well plates at a density of 2×10⁵ cells/well. After 24 hours of lactic acid treatment, washed with PBS. 1 ml of SA-β-gal staining working solution (prepared according to the instructions) was added to each well and incubated overnight (about 12-16 hours) in a 37°C CO₂-free incubator in the dark. After washing with PBS, observation and photography were performed under an optical microscope, and the proportion of SA-β-gal-positive (blue-stained) cells was counted.

### 10. Reactive oxygen species (ROS) level assay

Intracellular ROS was detected using an active oxygen detection kit (CA1410, Solarbio). CESCs were seeded in 6-well plates at a density of 2×10⁵ cells/well. After 24 hours of lactic acid treatment, the medium was removed. 2 ml of serum-free medium containing 10 μM DCFH-DA probe was added to each well and incubated at 37°C in the dark for 20 min. The probe solution was removed, and washed thoroughly with PBS 3 times. Flow cytometry detection: cells were digested with trypsin, collected, resuspended in PBS, and immediately detected for DCF fluorescence intensity (FL1 channel) on the machine. Fluorescence microscope observation: after washing with PBS, directly observe and take photos under a fluorescence microscope (IX 73, Olympus, Japan) (excitation wavelength 488 nm, emission wavelength 525 nm).

### 11. Immunofluorescence

Cells after treatment were fixed with 4% paraformaldehyde at room temperature for 15 minutes. Subsequently, they were blocked with bovine serum albumin (BSA) at room temperature for 1 hour, followed by incubation with primary antibody (γ-H2AX, KHC1551, Proteintech,1:200) at 4°C overnight. After that, the cells were incubated with secondary antibody (Dylight 488, Goat Anti-Rabbit IgG,A23220,Abbkine,1:1000) at room temperature in the dark for 1 hour. After rinsing with PBS, DAPI (1:1000 dilution) was added, and the cells were incubated at room temperature in the dark for 15 minutes. Finally, the experimental results were observed and imaged under a fluorescence microscope.

### 12. Western blotting

CESCs were lysed with RIPA lysis buffer (R0010, Solarbio) containing 1% PMSF (P0100, Solarbio) and lysed on ice for 30 min. The lysate was centrifuged at 12000×g for 10 min at 4°C, and the supernatant was collected. The protein concentration was determined using a BCA protein quantification kit (P0010S, Beyotime). An appropriate amount of protein sample was taken, 5×SDS-PAGE protein loading buffer (P1040, Solarbio) was added, and heated in a boiling water bath for 10 min. After the protein samples were separated by SDS-PAGE gel electrophoresis, the proteins were transferred to the PVDF membrane by wet transfer. The PVDF membrane was blocked with QuickBlock Western blocking solution (P0252, Beyotime) at room temperature for 1 hour. Add appropriately diluted primary antibody (P16, ab211542, Abcam,1:1000;P21, ab188224, Abcam,1:1000;P53, ab43615, Abcam,1:1000;γ-H2AX, ab81299, Abcam,1:1500) and incubate overnight at 4°C. After washing with TBST, add the corresponding HRP-labeled secondary antibody (HRP-conjugated Affinipure Rabbit Anti-Goat IgG(H+L), SA00001-4, proteintech,1:5000) and incubate at room temperature for 1 hour. After fully washing with TBST, develop using ECL chemiluminescence detection kit (1705060, Bio-Rad, USA), and collect images in ChemiDoc imaging system (Bio-Rad). Image J software was used to analyze the gray value of the target protein band.

### 13. Statistical analysis

All data are expressed as mean ± standard deviation (Mean ± SD). *In vitro* experiments were independently repeated at least 3 times (n=3), and *in vivo* experiments included at least 6 animals per group (n=6). Statistical analysis was performed using SPSS 21.0 and GraphPad Prism 8.0 software. Independent sample t-test (Student’s t-test) was used. *P* < 0.05 was considered statistically significant (\**P* < 0.05, \*\**P* < 0.01, \*\*\**P* < 0.001).

## Results

### 1. The puncture model confirms a positive correlation between lactic acid accumulation and IDD

A puncture-induced IDD model of the rat tail vertebra Co7/8 intervertebral space was successfully constructed (Fig 1A). X - ray showed that compared with the control group, the disc height index (DHI) in the puncture group was significantly reduced by 80.33% (*P*<0.001) (Fig 1B). Biochemical detection indicated that the lactic acid content in the IDD tissue reached (0.84 μmol/mg), which was significantly 7 times higher than that in the control group (0.12 μmol/mg) (*P*<0.001) (Fig 1C). HE and Safranin O-Fast Green (SO&FG) further confirmed that in the puncture group, the arrangement of nucleus pulposus cells was disordered, the staining intensity of proteoglycan was decreased, and the histological score was significantly increased (Fig 1D - E).

**Fig 1:**
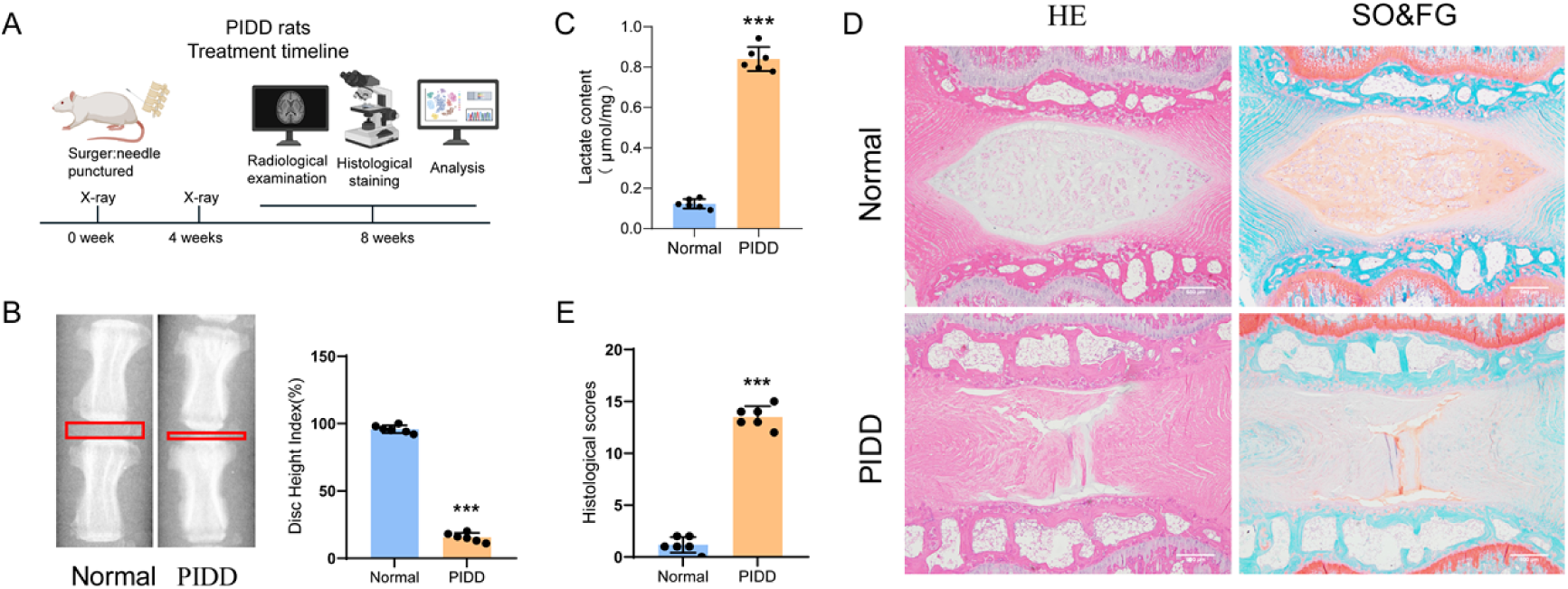
Accumulation of lactic acid in degenerated intervertebral discs: (A) Timeline of *in vivo* experimental design; (B) X-ray images of caudal vertebrae and intervertebral disc height index (IDHI) in normal and intervertebral disc-degenerated rats (n=6); (C) Determination of lactic acid content in intervertebral discs from normal rats and rats with painful intervertebral disc disease (PIDD) using a detection kit (n=6); (D-E) Hematoxylin-Eosin (HE) staining, Safranin O-Fast Green (SO&FG) staining, and histological scores of intervertebral discs in different groups.

### 2. Lactic acid inhibits CESC activity and induces oxidative stress

Following preliminary characterization, the isolated cartilage endplate stem cells exhibit stem cell characteristics (Fig S1 A-E). Upon treating CESCs with varying concentrations of lactic acid, CCK-8 assay results demonstrated a dose-dependent inhibition of cell viability. Specifically, after a 24-hour treatment with 2 mM, 6 mM, and 10 mM lactic acid, cell viability decreased to 65.5%, 58.4%, and 15.7% of the control group, respectively (all *P* < 0.001) (Fig 2A). Given that the inhibitory effect was most pronounced at the 24-hour time point, this treatment duration was selected for subsequent experiments. Both flow cytometry (using DCFH-DA) and fluorescence imaging techniques provided conclusive evidence that lactic acid treatment significantly elevated intracellular reactive oxygen species (ROS) levels. Flow cytometry data revealed that the average fluorescence intensities of the 2 mM, 6 mM, and 10 mM lactic acid treatment groups were 1.5-fold, 1.9-fold, and 3.8-fold higher than that of the control group, respectively (all *P* < 0.001) (Fig 2B). Similarly, fluorescence imaging showed that the average fluorescence intensities were 1.12-fold, 1.2-fold, and 1.5-fold greater than the control, respectively (all *P* < 0.001) (Fig 2C).

**Fig 2.**
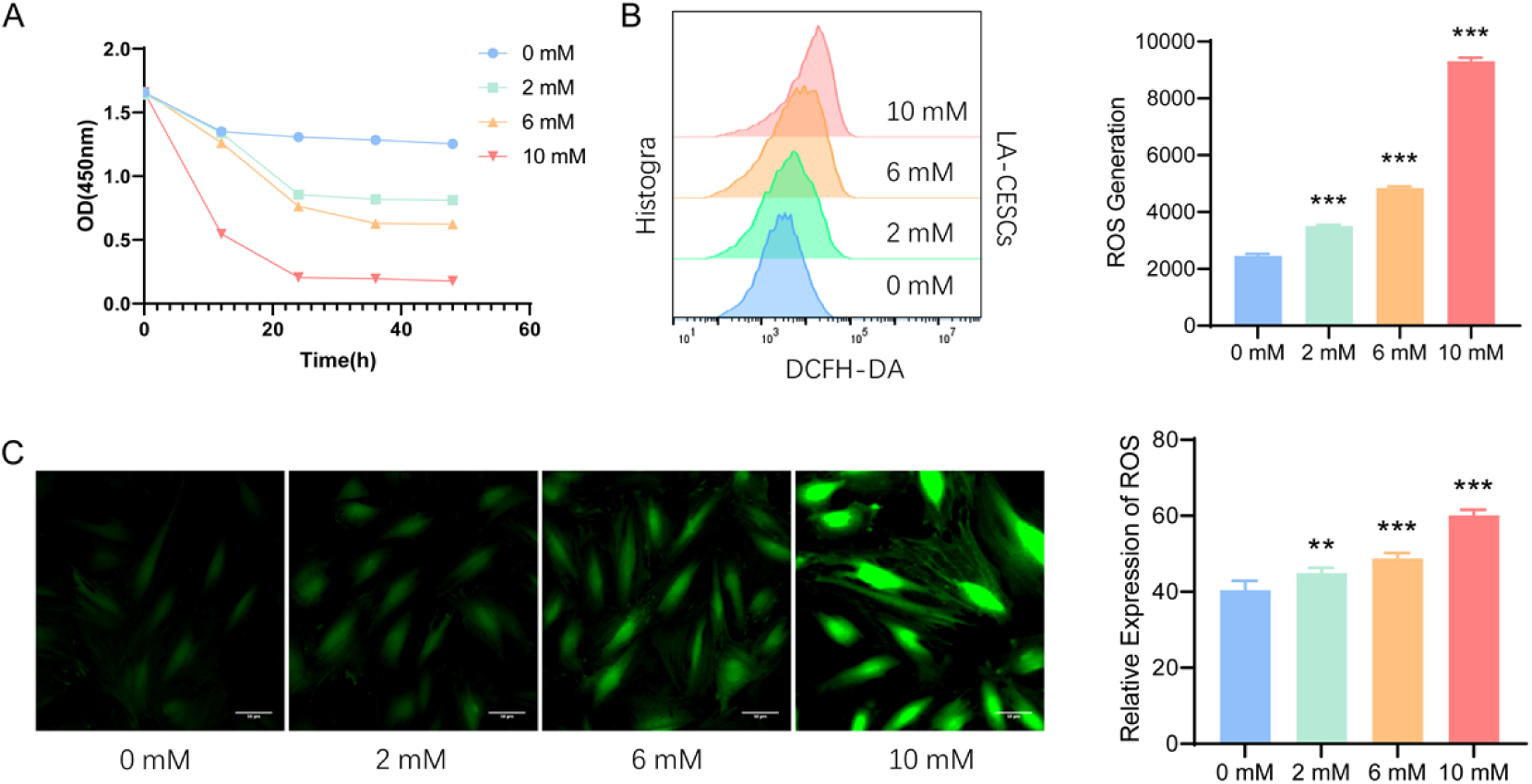
Lactic acid suppresses the activity of CESCs and triggers oxidative stress: (A) CCK-8 assay was used to evaluate the effects of lactic acid at different concentrations and treatment durations on CESCs viability; (B) Flow cytometry was performed to detect the effect of different lactic acid concentrations on intracellular reactive oxygen species (ROS) levels in CESCs; (C) Fluorescence-based detection assay was employed to analyze the effect of various lactic acid concentrations on ROS production in CESCs.

### 3. Lactic acid induces cell cycle arrest and proliferation inhibition of CESCs

Flow cytometry cell cycle analysis showed that lactic acid treatment significantly increased the proportion of cells in the G0 phase. Compared with the control group (38.16%), the proportion of cells in the G0 phase in the 2 mM, 6 mM, and 10 mM treatment groups increased to 53.03%, 69.67%, and 85.63%, respectively (*P*<0.001) (Fig 3A). EDU staining further confirmed that lactic acid inhibited DNA synthesis, and the EDU positive rate decreased from 23.5% in the control group to 12.35%, 11.01%, and 8.62%, respectively (*P*<0.001) (Fig 3B).

**Fig 3.**
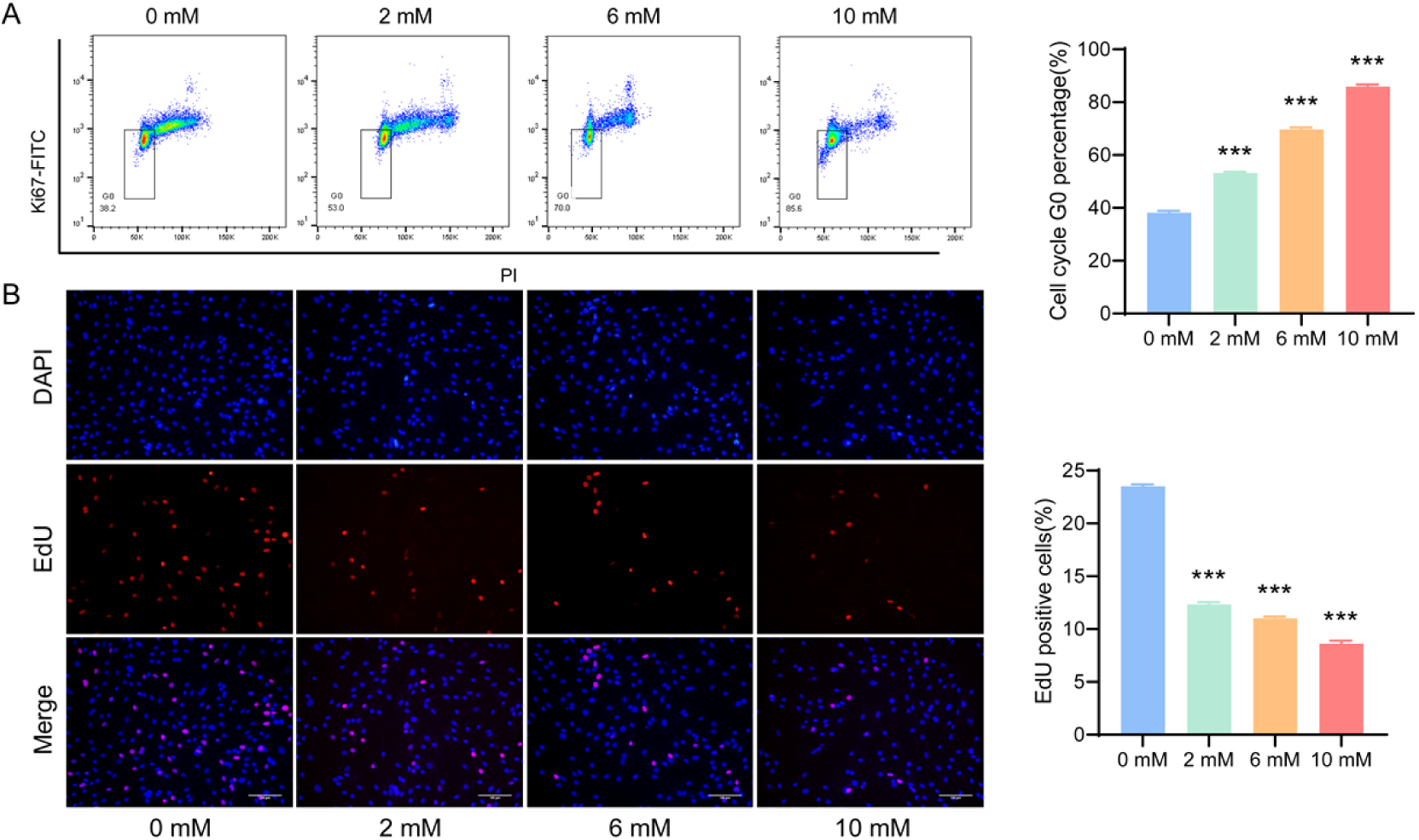
Lactic acid inhibits the proliferation of CESCs: (A) Effects of different concentrations of lactic acid treatment on the cell cycle distribution of CESCs; the boxed area indicates the G0 phase; (B) EdU incorporation assay was used to detect the effect of different concentrations of lactic acid treatment on CESCs proliferation. Proliferating CESCs are shown in red, and cell nuclei were stained with Hoechst 33342 (blue).

### 4. Lactic acid triggers senescence and DNA damage in CESCs

β - galactosidase staining showed that after 24 hours of lactic acid treatment, the proportion of positive cells increased significantly with the increase of concentration (0 mM: 8.89%; 2 mM: 23.06%; 6 mM: 27.24%; 10 mM: 33.06%, *P*<0.001) (Fig 4A - B). Immunofluorescence detection of the DNA damage marker γ - H2AX found that lactic acid treatment significantly increased the number of phosphorylated γ - H2AX foci (control group: 1.33 foci/cell; 2 mM group: 40.33 foci/cell; 6 mM group: 49.00 foci/cell; 10 mM group: 54.67 foci/cell, *P*<0.001) and fluorescence intensity (Fig 4C - D). Western blot analysis showed that lactic acid treatment significantly upregulated the expression of key regulatory proteins. In the 10 mM lactic acid treatment group, the expression levels of cell cycle inhibitory proteins P16 and P21 were 2.1 and 3.1 times that of the control group, respectively, the expression level of the pro - apoptotic/senescence factor P53 was 2.4 times that of the control group, and the expression level of the DNA damage marker γ - H2AX was 1.8 times that of the control group (*P*<0.001) (Fig 4G - I). The upregulation of protein expression was dose - dependent.

**Fig 4.**
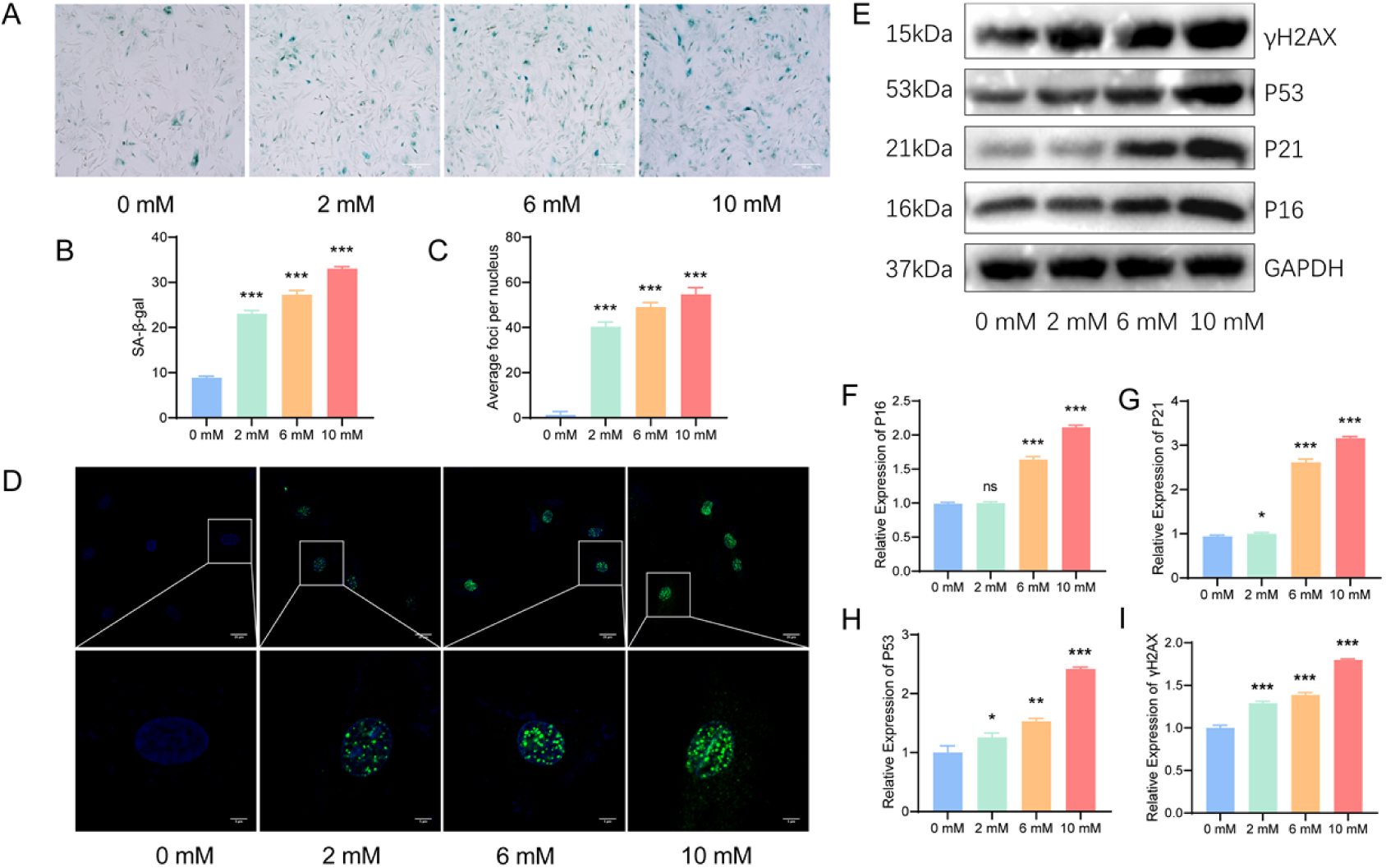
Lactic acid triggers the aging of CESCs and DNA damage: (A-B) Senescence-associated β-galactosidase (SA-β-gal) assay was performed to detect the effect of lactic acid treatment at different concentrations on CESCs senescence; senescent cells are visualized in blue; (C-D) γ-H2AX immunofluorescence assay was used to determine the effect of lactic acid treatment at different concentrations on DNA damage in CESCs; green foci indicate DNA damage sites, and cell nuclei were stained with DAPI (blue); (E-I)Western blotting was performed to assess the effects of different lactic acid concentrations on the expression levels of P16, P21, P53, and γ-H2AX proteins; the corresponding quantitative analysis results are also presented.

### 5. In vivo lactic acid injection directly exacerbates IDD

Following the intradiscal injection of gradient concentrations of lactic acid (0 mM, 6 mM, 10 mM) over an 8-week period, HE and SO&FG staining revealed notable differences. Compared to the control group, the high-concentration lactic acid group (10 mM) exhibited a substantial reduction in nucleus pulposus cell density and pronounced destruction of the matrix structure. Sirius red staining further demonstrated a disordered arrangement of collagen fibers (Fig 5A - B). X-ray analysis showed that the disc height index (DHI) values of the treatment groups were 95.12%, 46.67%, and 15.33% of the control group, respectively. Statistical analysis confirmed a significant dose-dependent decrease in DHI with lactic acid injection (*P*<0.001) (Fig 5C). These findings provide direct evidence that lactic acid promotes intervertebral disc degeneration in vivo.

**Fig 5.**
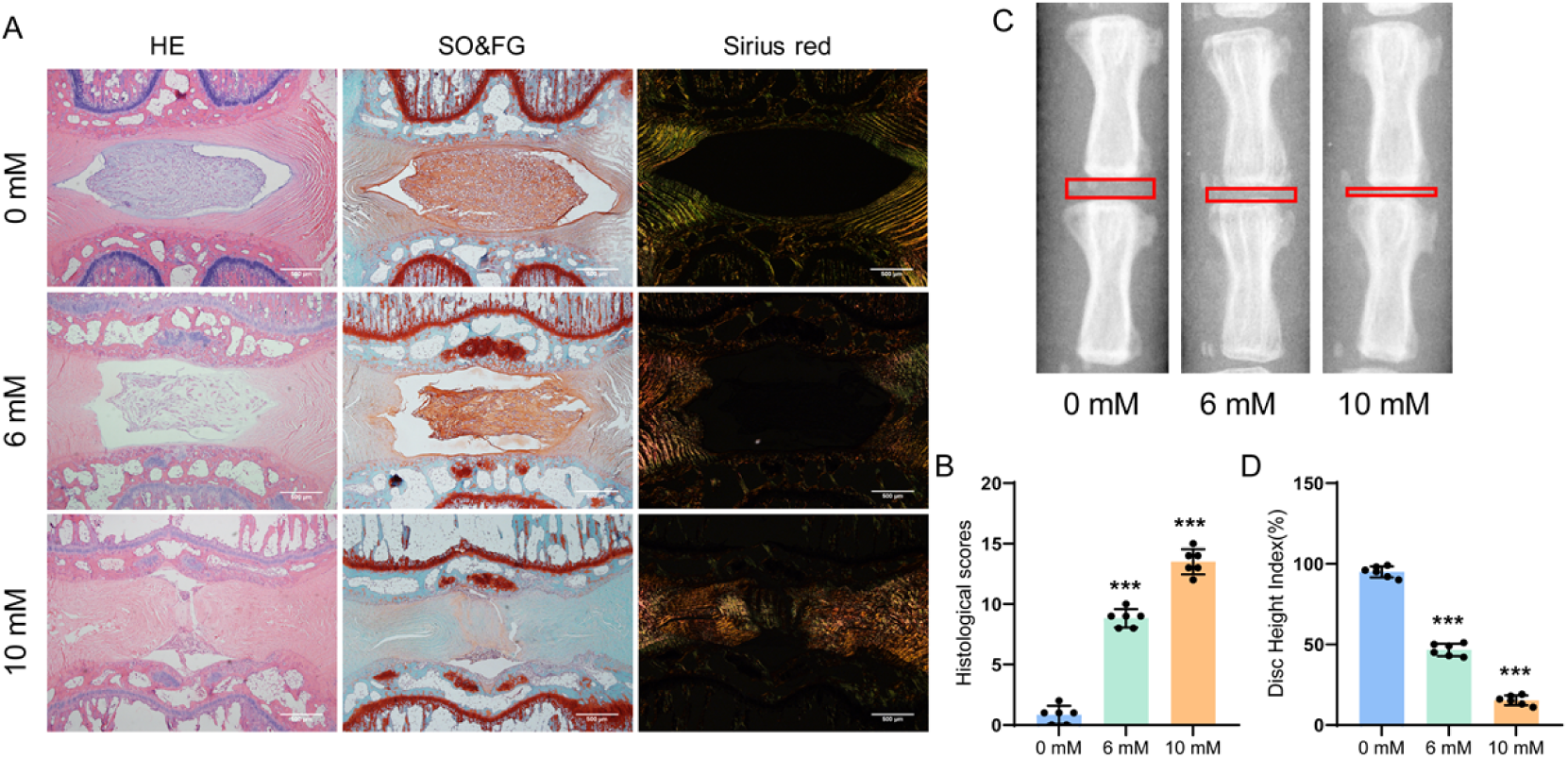
Intrathecal injection of lactic acid directly exacerbates intervertebral disc degeneration: (A-B) Hematoxylin-Eosin (HE), Safranin O-Fast Green, and Sirius Red staining of intervertebral discs following treatment with different concentrations of lactic acid, as well as the corresponding histological scores; (C-D) X-ray images of caudal vertebrae and intervertebral disc height index (IDHI) in specimens treated with different concentrations of lactic acid (n=6 per group).

**Fig 6.**
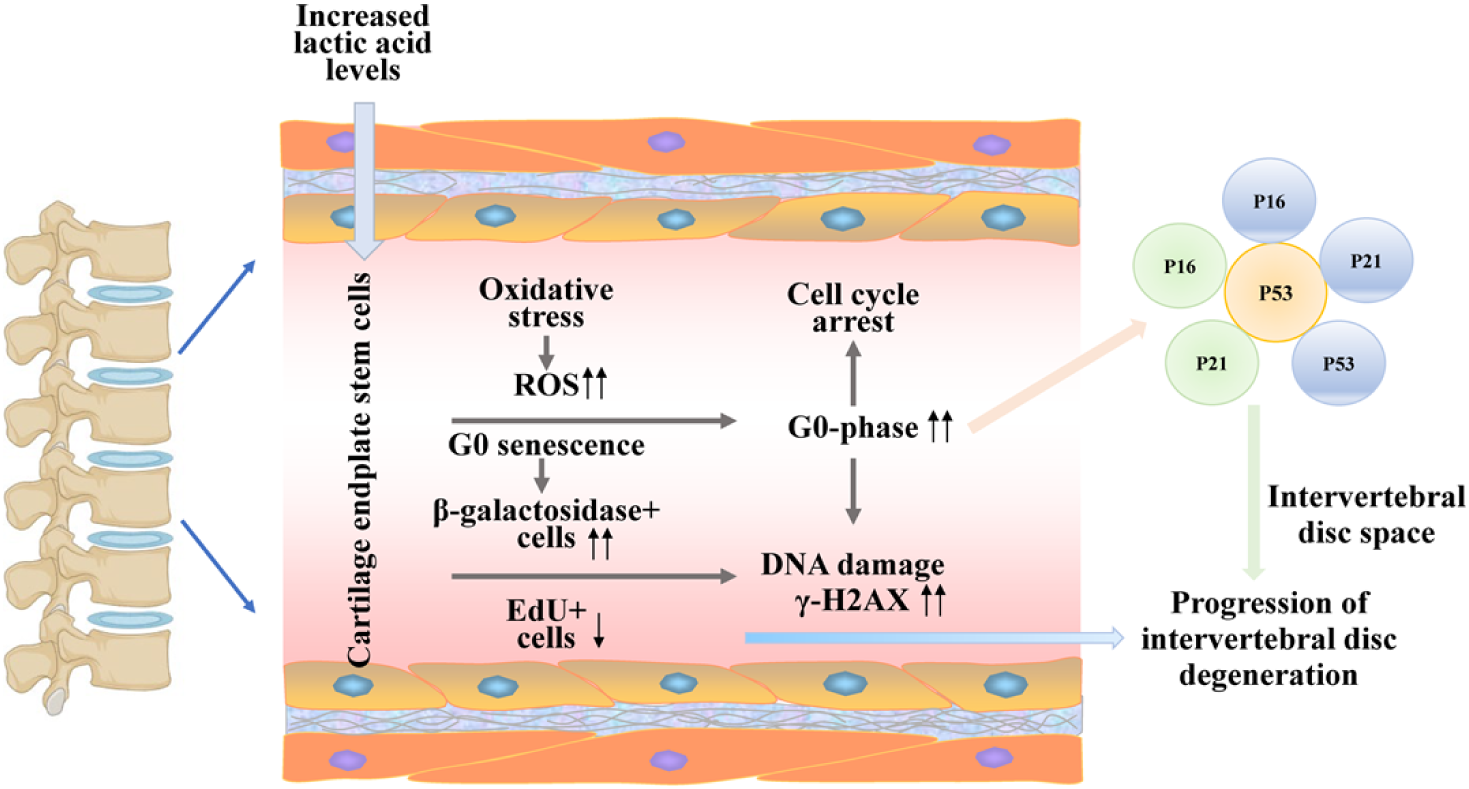
Mechanism of Lactic Acid-Promoted Intervertebral Disc Degeneration: Lactic acid promotes intervertebral disc degeneration by inducing oxidative stress, DNA damage, cell cycle arrest, and senescence in cartilage endplate stem cells (CESCs), thereby activating the signaling pathway centered on P16/P21/P53 and featuring γ-H2AX as a key molecule.

## Discussion

This study systematically elucidates, for the first time, the molecular mechanism of lactic acid: it induces senescence and DNA damage in CESCs, thereby activating the P16/P21/P53 pathway and ultimately driving IDD. This finding not only provides a novel perspective for deciphering the pathophysiological mechanisms of IDD but also lays a theoretical cornerstone for developing metabolism-targeted therapeutic strategies for IDD.

### 1. Pathological association between lactic acid accumulation and IDD

During the progression of intervertebral disc degeneration, functional cells such as nucleus pulposus cells and cartilage endplate cells are often exposed to a hypoxic microenvironment for long periods. As a result, they frequently rely on the glycolytic pathway for energy production—a metabolic pattern that ultimately leads to the accumulation of lactic acid, a metabolic byproduct^28,29^. In this study, using a rat caudal needle puncture-induced disc degeneration model, we found that lactic acid content in degenerated intervertebral discs was significantly increased, and its levels exhibited a positive correlation with both the degree of intervertebral space narrowing and the degree of nucleus pulposus structural destruction. Notably, this result is highly consistent with the findings confirmed by Li et al. via CEST-MRI technology—which clearly demonstrated a significant positive correlation between intervertebral disc lactic acid concentration and Pfirrmann grade (r=0.81, P<0.001). Notably, *in vitro* injection experiments further confirmed that lactic acid itself can exacerbate intervertebral disc degeneration—indicating it is not merely a passive byproduct of the degenerative process, but rather an active regulatory factor that drives the occurrence and development of intervertebral disc degeneration (IDD)^4^. This finding provides direct experimental evidence for the academic hypothesis that “metabolic reprogramming is a core driver of IDD” ^30^.

### 2. Damaging effects of lactic acid on the functions of CESCs

As seed cells pivotal for cartilage endplate repair, the preservation of both the proliferative capacity and differentiation potential of CESCs is fundamental to maintaining intervertebral disc homeostasis^31,32^. This investigation demonstrated that lactic acid exerted an inhibitory effect on CESC activity in a dose-dependent fashion. Specifically, exposure to 10 mM lactic acid led to a significant reduction in cell viability, decreasing it to 15.7% of the control group levels. Concurrently, the EDU positive rate plummeted to 8.62%, while 85.63% of cells were arrested at the G0 phase of the cell cycle. These findings align with the results reported by Liu et al. (2025) in a 3D organoid model, where 10 mM lactic acid was shown to impede the chondrogenic differentiation of CESCs by down-regulating the expression of SOX9 and COL2A1 by 60-70%^28^. Subsequent analyses revealed that lactic acid treatment significantly increased the proportion of β-galactosidase positive cells in CESCs to 33.06%, indicative of cellular senescence. Collectively, these results suggest that lactic acid may deplete the CESC reservoir, compromise the self-repair capacity of the cartilage endplate, and ultimately contribute to the progression of intervertebral disc degeneration^30^.

### 3. Cascade reaction of oxidative stress and DNA damage

This study found that lactic acid-induced reactive oxygen species (ROS) accumulation—with ROS levels in the 10 mM lactic acid treatment group being 2.8-fold higher than those in the control group—is a key upstream event that triggers damage to cartilage endplate stem cells. A As powerful oxidants, reactive oxygen species (ROS) can directly attack DNA double strands and thereby induce the formation of characteristic γ-H2AX foci following DNA double-strand damage^33–35^. This finding complement those of Li et al. (2025) at the mechanistic level: they demonstrated that an acidic microenvironment (pH 6.6) mediates ROS burst in a NOX4-dependent manner—a process that not only induces γ-H2AX foci formation in cartilage endplate stem cells (CESCs) (showing a 3.2-fold increase compared to the control group) but also promotes phosphorylation of the ATM/CHK2 signaling pathway^36^. Additionally, the expression level of γ-H2AX in the lactic acid treatment group was 1.8-fold higher than that in the control group—a result that further confirms the occurrence of intervertebral disc degeneration-related DNA damage. Previous studies have confirmed that persistent DNA damage can activate the P53/P21 pathway, thereby inducing permanent cell cycle arrest ^33,37–39^. In the present study, we observed a significant increase in the levels of P53 (with a 2.4-fold upregulation in expression) and P21 (with a 3.1-fold upregulation in expression)—a finding that serves as direct evidence of the aforementioned mechanism in our study system.

### 4. Core role of the P16/P21/P53 pathway in lactic acid-induced senescence

In the molecular regulatory network of cellular senescence, the P16/Rb and P53/P21 pathways are the two core signal axes ^24,40–42^. This study found that lactic acid can simultaneously up-regulate the expression of P16 (2.1-fold), P21 (3.1-fold) and P53 (2.4-fold), suggesting that it may accelerate CESC senescence by synergistically activating the two pathways. This finding is mutually verified with the gene intervention experiment of Balaraman et al. (2025): targeted silencing of p16 (siRNA) can reduce SA-β-gal⁺ cells by 58% in CESCs treated with lactic acid, while p21 knockout reverses G0 phase arrest (G0 proportion ↓44%)^43^. It is worth noting that as the core molecule of DNA damage response, the activation of P53 can not only induce cell cycle arrest through P21 but also directly initiate the apoptosis program^40,42,44^, which provides a molecular basis for explaining how lactic acid mediates CESC senescence and functional loss simultaneously.

### 5. Clinical significance and limitations

The “lactic acid - oxidative stress - cell cycle arrest - cellular senescence” axis elucidated in this study offers a novel intervention target for intervertebral disc degeneration treatment. Potential therapeutic strategies may include targeting lactic acid transport with inhibitors such as MCT1 inhibitor AZD3965^45–47^, enhancing lactic acid clearance through nanoparticles loaded with lactate oxidase^47^, or directly intervening in the cellular senescence pathway using agents like the p16/p21 inhibitor dasatinib/quercetin^48^. However, several limitations should be noted:(1) Although the lactic acid concentration (10 mM) employed in *in vitro* experiments aligns with the microenvironment of degenerated intervertebral discs^49^, the dynamic metabolic processes occurring in vivo are far more intricate;(2) The study did not explore the regulatory effects of lactylation modifications, such as H3K18la and H4K12la, on p16/p21 transcription. Emerging evidence from recent studies indicates that histone lactylation can directly activate senescence-associated genes^28,50^;(3) Rescue experiments involving the specific knockout of P16/P21/P53 to reverse lactic acid-induced IDD were absent. Future research should integrate gene-edited animal models and clinical samples to further validate the pathophysiological significance of this proposed mechanism.

### Conclusion

This study systematically and explicitly confirms, for the first time, that lactic acid is a core metabolic factor driving the occurrence and progression of intervertebral disc degeneration. This factor induces sequentially oxidative stress bursts, DNA damage, cell cycle arrest, and senescence in cartilage endplate stem cells, and thereby activates the signaling pathway centered on P16/P21/P53 with γ-H2AX as the key molecule. This ultimately forms an “oxidative stress-cell cycle arrest-cell senescence regulatory axis,” disrupting intradiscal homeostasis and propelling the progression of IDD. These findings at the mechanistic level not only provide critical new perspectives and experimental support for the in-depth interpretation of the pathophysiological mechanisms of IDD but also, more importantly in terms of translational value, define clear core directions and novel avenues for developing innovative preventive and therapeutic strategies for IDD—by targeting the regulation of the lactic acid-CESC senescence axis.

## Acknowledge

We would like to thank all participants for their advice and support in this study.

## Data availability statement

Data will be made available on request.

## Author Contributions

Qiao Lv and Tianling Wang contributed equally to this work. Qiao Lv designed the research framework, conducted key experiments, analyzed primary data, and drafted the manuscript. Tianling Wang carried out supplementary experiments, validated data, and revised the manuscript. Lu Jiang, Qian Chen, Jin Peng, Ju Zhou, and Qian Min assisted with experiments, sample collection, data sorting, and literature review, offering technical support and preliminary analysis. Yongqi Pu provided advice on experimental methods, assisted in data interpretation, and optimized the research design.Jianyun Zhou and Qing Huang jointly supervised the project, conceived the research idea, guided the study direction, and finalized the manuscript. Both are responsible for the work’s integrity and correspondences. All authors reviewed and approved the submitted manuscript.

